# IFNγ and iNOS-mediated alterations in the bone marrow and thymus and its impact on *Mycobacterium avium*-induced thymic atrophy

**DOI:** 10.1101/2021.02.23.432464

**Authors:** Palmira Barreira-Silva, Rita Melo-Miranda, Claudia Nobrega, Susana Roque, Cláudia Serre-Miranda, Margarida Borges, Daniela de Sá Calçada, Samuel M. Behar, Rui Appelberg, Margarida Correia-Neves

**Affiliations:** Life and Health Sciences Research Institute (ICVS), School of Medicine, University of Minho, Braga, Portugal.; ICVS/3B’s, PT Government Associate Laboratory, Braga/Guimarães, Portugal.; Institute for Biomedicine (iBiMED), Department of Medical Sciences, University of Aveiro, Aveiro, Portugal.; UCIBIO/REQUIMTE, Departamento de Ciências Biológicas, Faculdade de Farmácia, Universidade do Porto, Portugal.; Department of Microbiology and Physiological Systems, University of Massachusetts Medical School, Worcester, Massachusetts, USA.; i3S, Instituto de Investigação e Inovação em Saúde, Universidade do Porto, Portugal.; IBMC-Instituto de Biologia Molecular e Celular, Universidade do Porto, Porto, Portugal.; ICBAS-Instituto de Ciências Biomédicas Abel Salazar, Universidade do Porto, Porto, Portugal.

## Abstract

Disseminated infection with the high virulence strain of *Mycobacterium avium* 25291 lead to progressive thymic atrophy. We previously uncovered that *M. avium*-induced thymic atrophy is due to increased levels of glucocorticoids synergizing with nitric oxide (NO) produced by interferon gamma (IFNγ) activated macrophages. Where and how these mediators are playing, was yet to be understood. We hypothesized that IFNγ and NO might be affecting bone marrow (BM) T cell precursors and/or T cell differentiation in the thymus. We show that *M. avium* infection causes a reduction on the percentage of lymphoid-primed multipotent progenitors (LMPP) and common lymphoid progenitors (CLP). Additionally, BM precursors from infected mice are unable to reconstitute thymi of RAGKO mice in an IFNγ-dependent way. Thymi from infected mice presents a NO-dependent inflammation. When transplanted under the kidney capsule of non-infected mice, thymic stroma from infected mice is unable to sustain T cell differentiation. Finally, we observed increased thymocyte death via apoptosis after infection, independent of both IFNγ and iNOS, and a decrease on activated caspase-3 positive thymocytes, that was not observed in the absence of iNOS expression. Together our data suggests that *M. avium*-induced thymic atrophy results from a combination of impairments, mediated by IFNγ and NO, affecting different steps of T cell differentiation from T cell precursor cells in the BM to the thymic stroma and thymocytes.

## INTRODUCTION

T cell mediated immunity is crucial during the immune response to mycobacteria infections. T cells differentiate within the thymus, from progenitors that migrate from the bone marrow (BM). In the thymus, T cell precursors find the specialized environment required for their differentiation into new T cells. The thymus undergoes physiological involution with aging ^1,2^. This fact, together with reports from late 1990s showing that people whose thymus was removed at early age seemed to present a normal T cell repertoire when adults, led to the idea that the thymus was only needed for production of T cells early in life ^3^. However, further investigation clearly showed that people thymectomized early in life develop premature immune aging ^4^. In addition, the amount of functional thymic tissue correlates with the reconstitution of the peripheral T cell repertoire in cases of immune-reconstitution after severe lymphopenia, as is the case of patients with acquired immunodeficiency syndrome (AIDS) under antiretroviral therapy, or in cancer patients after BM transplant ^5,6^. These data raised the relevance of understanding premature thymic atrophy since it can impact on the overall immune health of the individual.

A variety of pathogens target the thymus and induce premature thymic atrophy both in humans and experimental animal models, such as the bacterium *Mycobacterium avium*, the fungus *Paracoccidioids brasiliensis*, the parasite *Tripanosoma cruzi*, and the virus HIV, among others ^7,8^. Although the thymus is a target for infection with different mycobacteria ^9^, *M. avium*-induced thymic atrophy has been shown to be strain-dependent ^10^. Our studies revealed premature thymic atrophy after infection with the high virulence *M. avium* strain 25291, but no signs of atrophy with the low virulence strain 2447, even after several months of chronic infection ^10,11^. *M. avium* infection-induced thymic atrophy arises from a synergy between glucocorticoids (GC) and nitric oxide (NO) produced by interferon gamma (IFNγ) activated macrophages (Mϕ)^10^. However, where and the mechanism(s) by which these mediators act to lead to *M. avium* infection-induced thymic atrophy is unknown. Here we hypothesized that mechanisms mediated by IFNγ and inducible NO synthase (iNOS) may impact the T cell differentiation in the thymus and/or earlier, the BM T cell precursor cells, resulting in the development of *M. avium*-induced thymic atrophy.

Several research groups described different alterations in the thymus, underlying premature thymic atrophy, using diverse mouse infection models ^7,8,12^. Common mechanisms are: (1) thymic structural changes ^13,14^; (2) reduction in thymocyte proliferation ^15^; (3) increased thymocyte cell death ^16–19^; and, (4) increased export of immature thymocytes to the periphery ^20–22^. One of the most described mechanisms, is that GC overproduction lead to excessive death of thymocytes, mostly at the double positive (DP) stage ^23,24^. Additionally, it is known that IFNγ and NO can also induce thymocyte cell death, either independently or by synergizing with GC ^16,25–28^.

Mechanisms responsible for premature thymic atrophy involving alterations of T cell precursors in the BM are seldom explored, and include: (1) decreased production; (2) arrest in the BM and consequent decreased migration and/or (3) loss of ability to settle the thymus. On the other hand, infection-induced alterations of the BM have been abundantly documented following infection by bacteria including *Mycobacterium tuberculosis* ^29^, *M. avium* ^30^, Group A *Streptococcus* ^31^ and *Escheriachia coli* ^32^; viruses including the Human Immunodeficiency Virus (HIV) ^33^, Pneumovirus ^34^ and Vaccinia virus ^35^; and parasites such as *Plasmodium chabaudi* ^36^, *Plasmodium berghei* ^37^ and *Trypanosoma brucei* ^38^. IFNγ is a major mediator that promotes BM cell alterations during some of these infections ^30,31,34,36^, via the expansion of LSK (Lineage^−^ Sca1^+^ cKit^+^) cells and the increased differentiation of cells from the myeloid lineage. Other molecules such as tumor necrosis factor (TNF) and NO have also been associated with BM alterations during infection ^34,38^. However, except for a report showing that the reduction of BM precursors is associated with sepsis-induced thymic atrophy ^39^, to our knowledge, no other reports associate BM alterations with infection-induced thymic atrophy. We previously showed that *M. avium* strain 25291 infected WT mice present a reduction in the number of early thymic precursors (ETP) ^10^, which is the most immature cell population in the thymus. Additionally, mice that lack iNOS expression (iNOSKO mice), known to be resistant to infection-induced premature thymic atrophy, also present reduced numbers of ETP^10^. Altogether, these data indicate that alterations in the BM T cell precursors, although not sufficient to cause premature thymic atrophy, might be an important step in the process. Here we took advantage of KO mouse strains, and of cell transfer and thymic transplant models to study the mechanisms mediated by IFNγ and/or iNOS affect the thymus and/or the BM leading to premature thymic atrophy during *M. avium* infection.

## METHODS

### Mice and infection

C57BL/6 wild-type (WT) mice were purchased from Charles River Laboratories (Barcelona, Spain) or bred at the Life and Health Sciences Research Institute (ICVS - School of Medicine, University of Minho, Braga, Portugal) from a breeding pair purchased from Charles River Laboratories. B6.SJL-*Ptprca Pepcb*/BoyJ (WT CD45.1) and IFNγ–deficient C57BL/6 (IFNγKO) mice were bred at ICVS from a breeding pair purchased from The Jackson Laboratory (Bar Harbor, ME, USA). Transgenic mice with a selective impairment on IFNγ signaling in CD68^+^ cells (MIIG) ^40^ were bred at the Institute for Molecular and Cell Biology (University of Porto, Porto, Portugal) from a breeding pair provided by the Cincinnati Children’s Hospital Medical Center and the University of Cincinnati College of Medicine (Cincinnati, OH, USA). iNOS-deficient C57BL/6 (iNOSKO) ^41^ mice were bred at ICVS after back-crossing the original strain (kindly provided by Drs. J. Mudgett, J.D. MacMicking, and C. Nathan, Cornell University, New York, NY, USA) onto a C57BL/6 background for seven generations, or purchased from The Jackson Laboratory. B6(Cg)-*Rag2^tm1.^*^1Cgn^ (RAGKO) mice were bred at ICVS from a breeding pair purchased from Instituto Gulbenkien da Ciência (Oeiras, Portugal). iNOS and Rag2-double deficient (iNOS.RAG.2KO) mice were obtained, at ICVS, after crossing F1 resulting from the cross of iNOSKO and RAGKO mice.

Eight- to ten-week-old female mice were infected intravenously (i.v.) through a lateral tail vein with 10^6^ CFU of the *M. avium* strain ATCC 25291 SmT (obtained from the American Type Culture Collection, Manassas, VA, USA) or the strain 2447 (provided by Dr. F. Portaels, Institute of Tropical Medicine, Antwerp, Belgium). Bacterial inocula preparation and bacterial load quantification in the organs was performed as previously described ^11,42^. Although no signs of major distress were observed for the first 2 months upon infection, some animals showed signs of deterioration of body condition in a non-synchronous way. To avoid excessive and unnecessary suffering of animals, humane endpoints were applied and mice were euthanized when reaching 25% weight loss.

Mice euthanasia was performed through controlled CO_2_ inhalation or an overdose of Ketamine (150 mg/kg) + Medetomidine (2 mg/kg) injected intraperitoneally (i.p.), followed by lethal blood collection (performed after confirmation of anaesthesia) and thoracotomy.

All experiments were performed in accordance with the recommendations of the European Convention for the Protection of Vertebrate Animals Used for Experimental and Other Scientific Purposes (ETS 123) and 86/609/EEC Directive and Portuguese rules (DL 129/92). The animal facility and people directly involved in animal experiments were certified by the Portuguese regulatory entity - *Direção Geral de Alimentação e Veterinária* (DGAV); the animal experimental protocols were approved by DGAV (# 015584), or by the Institutional Animal Care and Use Committee at the University of Massachusetts Medical School (Animal Welfare A3306-01), using the recommendations from the Guide for the Care and Use of Laboratory Animals of the National Institutes of Health and the Office of Laboratory Animal Welfare.

### Single cell suspensions

Spleens, thymi and BM were collected and processed individually. Single cell suspensions were obtained from thymi and spleens by gentle mechanical dissociation in complete DMEM (cDMEM - DMEM supplemented with 10% heat-inactivated FCS, 10 mM HEPES, 1 mM sodium pyruvate, 2 mM L-glutamine, 50 mg/ml streptomycin, and 50 U/ml penicillin (all from Invitrogen Life Technologies); and from femurs by gentle flush of the BM with cDMEM. RBCs were lysed using a hemolytic solution (155 mM NH_4_Cl, 10 mM KHCO_3_, pH 7.2) for 4 min at room temperature, and cells were re-suspended in cDMEM. The number of viable cells was counted by trypan blue exclusion using a hemocytometer.

### Flow cytometry

One million cells were stained for flow cytometry analysis. For BM analysis, cells were stained with FITC-conjugated anti-lineage markers [anti-CD3 (145-2C11), anti-CD4 (RM4-5), anti-CD8 (53-6.7), anti-CD19 (6D5), anti-B220 (RA3-6B2), anti-CD11b (M1/70), anti-CD11c (N418), anti-NK1.1 (PK136), anti-Gr1 (RB6-8C5), anti-TER119 (TER119)], PE-conjugated anti-cKit (2B8), PerCP-Cy5.5-conjugated anti-Sca1 (D7), PE-Cy7–conjugated anti-IL7Rα (A7R34), APC-conjugated anti-Flt3 (A2F10), APC-Cy7-conjugated anti-CD48 (HM48-1) and BV421-conjugated anti-CD150 (TC15-12F12.2). For thymocyte analysis, cells were labeled with distinct combinations of the following antibodies: FITC, PerCP-Cy5.5, BV510 or V500-conjugated anti-CD8 (53-6.7), PerCP-Cy5.5-conjugated anti-CD24 (M1/69), Pe or APC-conjugated anti-CD3 (145-2C11), Pe-Cy7 or APC-Cy7-conjugated anti-CD44 (IM7), APC-Cy7 or V450-conjugated anti-CD4 (RM4-5), Alexa647-conjugated anti-active caspase3 (C92-605) and APC-conjugated Annexin V. For live/dead cell analysis, propidium iodide (Pi; Sigma-Aldrich, Germany) was added at a final concentration of 1 mg/ml, 15 min before acquisition on the flow cytometer. For spleenocyte analysis, cells were labeled with the following antibodies: FITC-conjugated anti-CD11b (M1/70), APC-conjugated anti-CD3 (145-2C11), APC-Cy7-conjugated anti-CD19 (6D5), V450-conjugated anti-CD4 (RM4-5) and V500-conjugated anti-CD8 (53-6.7). All antibodies were purchased from BioLegend (San Diego, CA, USA) except the V450-conjugated anti-CD4 (RM4-5), the V500-conjugated anti-CD8 (53-6.7) and the Alexa647-conjugated anti-active caspase3 (C92-605), which were obtained from BD Biosciences (San Jose, CA, USA). Acquisition was performed on a LSRII flow cytometer using BD FACSDiva software v6.0 (Becton and Dickinson, NJ, USA), or on a MACSQuant flow cytometer (Miltenyi Biotec, Germany). Data were analyzed using FlowJo 10.7.1 (BD Biosciences).

### Thymic transplant

Thymi were aseptically removed from non-infected RAGKO or from WT mice infected for 70 days with *M. avium* 25291. Thymi were maintained in cDMEM for no longer than 20 min until being transplanted under the kidney capsule of WT CD45.1 mice (anesthetized with 200mg xylazine hydrochloride and 200mg ketamine hydrochloride, administered i.v.). One thymic lobe from non-infected RAGKO mice and one thymic lobe from infected WT mice were transplanted to the same WT CD45.1 mouse (one on each kidney). Mice were euthanized 4 weeks after transplant as described earlier, and the transplanted thymi analyzed by flow cytometry.

### BM adoptive transfer

Single-cell suspensions of pools of BM cells were prepared from non-infected or from 70 days *M. avium* 25291 infected WT or IFNγKO mice. BM progenitor cells were purified from each suspension using the Lineage Cell Depletion Mouse Kit microbeads (Miltenyi Biotec). Magnetic separation was performed with an autoMACS separator (Miltenyi Biotec). After purification, viable cells were counted by trypan blue exclusion using a hemocytometer; purity was confirmed by flow cytometry stain using FITC-conjugated anti-lineage markers and PE-conjugated anti-cKit. Cells (1-1.5 × 10^6^) were transferred i.v. to each RAGKO or iNOS.RAG.2KO mouse treated with Busulfan (0.6mg/mouse) i.p. 24 hours before. Recipient mice received prophylactic antibiotic treatment *ad libitum* [2,5% of Bactrim (40 mg trimethoprim + 200 mg sulfametoxazol) in drinking water] for 5 days (treatment finished 2 days before BM transference). Mice were euthanized 4 weeks after cell transfer and the recipient thymus and spleen were analyzed by flow cytometry.

### Measurement of corticosterone serum levels

To obtain the serum concentration of corticosterone at basal levels, blood samples were collected at 9 am (1 h after lights on) from a venous incision at the tip of the tail during a period not exceeding 2 min for each mouse (to avoid corticosterone sera level increase due to handling). Sera were isolated by centrifugation and stored at −80°C. Corticosterone levels were determined using Corticosterone ELISA kit (ENZO life sciences, Inc., NY, USA) according with manufacturer’s instructions.

### Real Time-quantitative PCR analysis

Total RNA was extracted from 1 thymic lobe (except for 70 dpi, where the whole thymus was used due to severe atrophy) using TRIzol™ Reagent (Invitrogen, Carlsbad, CA, USA). RNA was quantified using NanoDrop 2000 (Thermo Scientific, MA, USA), and 1 μg was run on a 1% agarose gel to check for RNA integrity. Complementary DNA (cDNA) was synthesized using iScriptTM Advanced cDNA Synthesis Kit (Bio-Rad Laboratories, CA, USA), according to manufacturer’s instructions. Real Time-quantitative PCR (RT-qPCR) was performed using the SsoFast EvaGreen Supermix (Bio-Rad Laboratories, CA, USA) and the primer pairs described in Table 1. Expression of target and housekeeping genes (see Table 1) was analysed on CFXTM Manager (Bio-Rad Laboratories) using the “Gene Study” function, and exported to Microsoft Excel for further calculations.

**Table 1.**
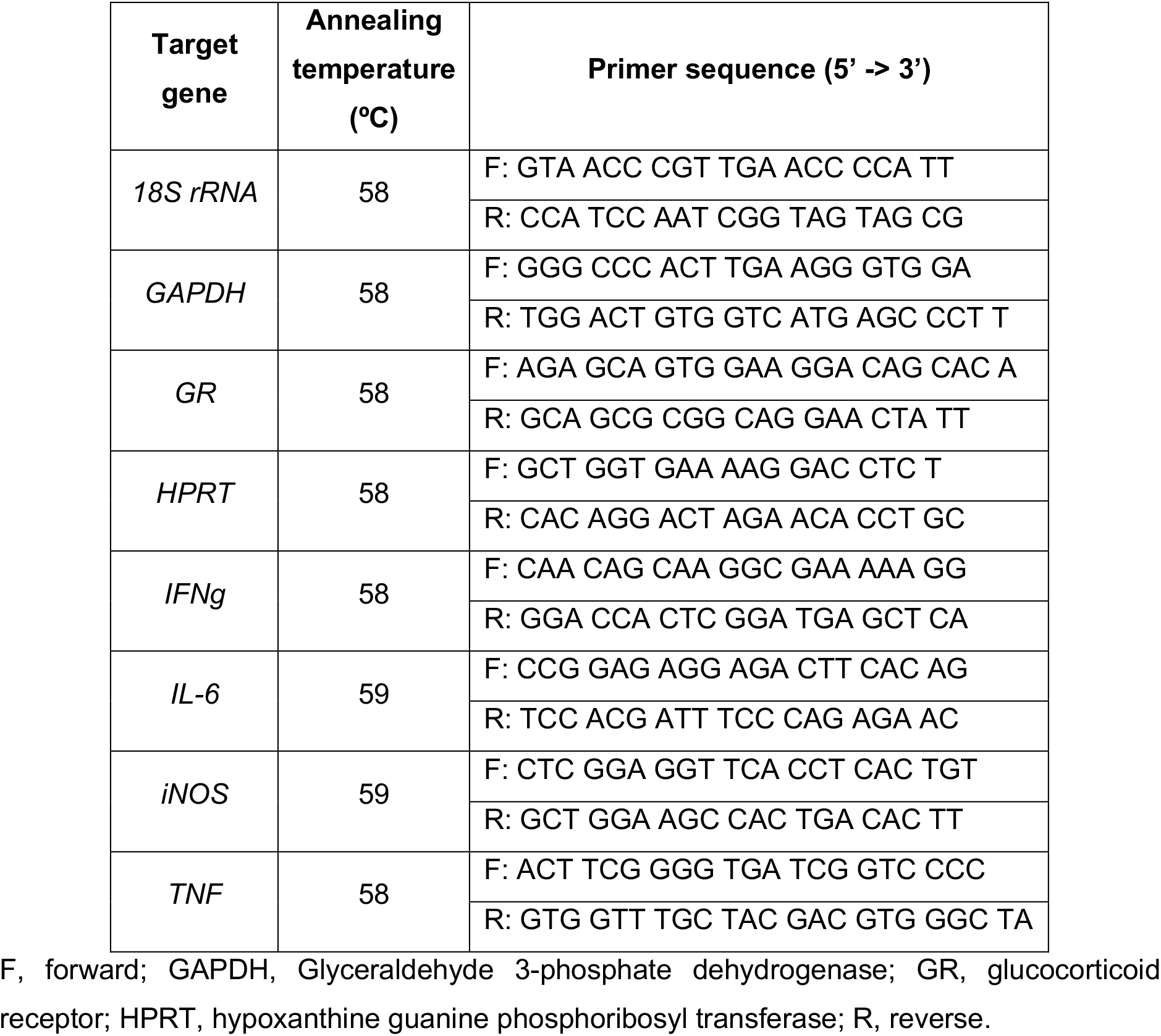
Genes analysed by RT-qPCR. Primer sequences and respective annealing temperatures.

### Statistical analysis

Results were expressed as mean or mean + SD, unless otherwise said. Prism 8.4.3 (GraphPad) was used for all the statistical analysis. Variables normality was accessed by Kolmogorov-Smirnov test. Parametric tests were used (as indicated on figure legends): two-tailed unpaired *t*-test; two-tailed ratio paired *t*-test; ordinary one-way ANOVA followed by Dunnett’s multiple comparisons test; 2-way ANOVA followed by Tukey’s multiple comparisons test. Differences between groups were considered statistically significant for *p-*values <0.05.

## RESULTS

### Exacerbated proportion of LSK cells in the BM of mice infected with *M. avium* strain 25291 is dependent on IFNγ but not iNOS

The differentiation of new T cells is dependent on the continuous seeding of BM T cell precursors in the thymus. To identify the alterations caused by *M. avium* infection among T cell precursors within the BM, potentially associated with infection-induced thymic atrophy, we compared the BM from wild type (WT) mice infected with *M. avium* strain 25291 (that present premature thymic atrophy) with the BM from WT mice infected with *M. avium* strain 2447 (that does not present premature thymic atrophy). Additionally, we analyzed the BM from mice lacking the expression of two major players of *M. avium* strain 25291 infection-induced thymic atrophy, IFNγ and iNOS, or the signaling of IFNγ in Mϕ (the three mouse strains - IFNγKO, iNOSKO and MIIG - are protected from premature thymic atrophy).

We observed that the bacterial load in the BM from mice infected with the strain 25291 was greater compared to that of mice infected with strain 2447, reaching a 5-log difference at 70 days post-infection (dpi; Fig. 1A). At 80dpi with *M. avium* strain 25291, mice lacking IFNγ had similarly high bacterial load in their BM compared to WT mice (Fig. 1B) and iNOSKO mice showed 2-log lower bacterial load than WT mice (Fig. 1C), in accordance with what was previously described regarding bacterial load in other organs such as the liver, spleen, lung and thymus ^10,11^. Infection of WT mice with strain 25291 led to decreased number of total BM cells (Fig. 1D), as previously described ^43^. On the other hand, the total number of BM cells from WT mice infected with strain 2447 did not differ from those of non-infected animals up to 80 dpi (Fig. 1D).

**Figure 1.**
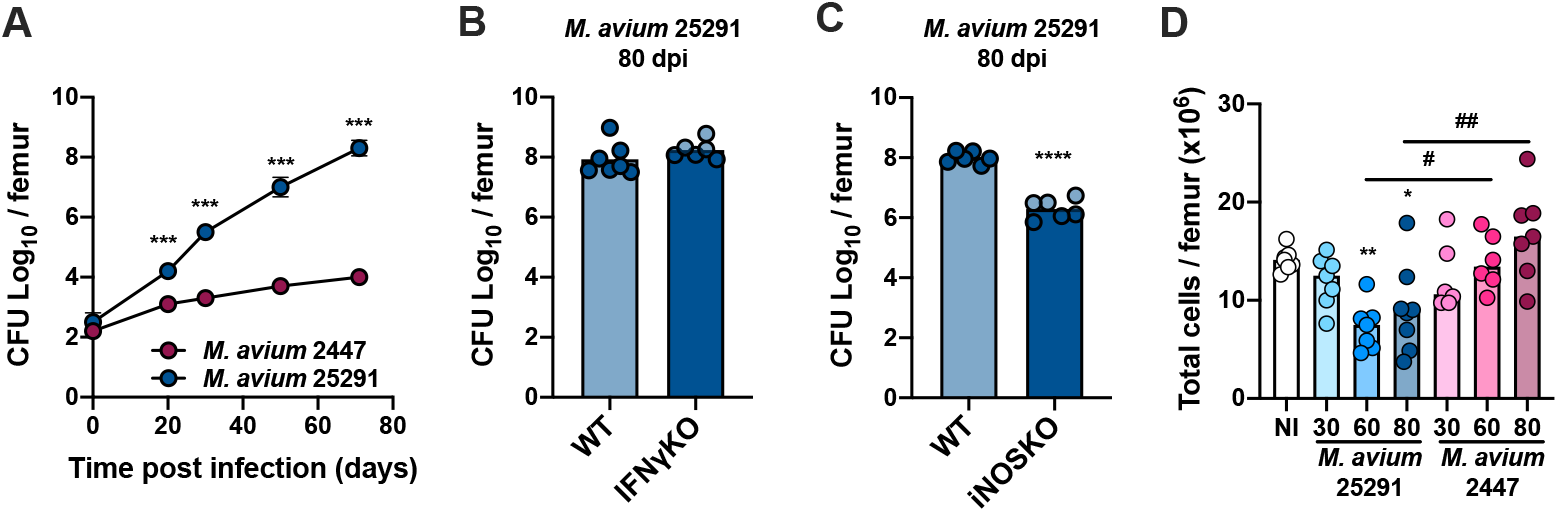
Bacteria load in the BM is higher after infection with *M. avium* strain 25291 than with strain 2447. (A) Representative kinetics of *M. avium* infection in the BM of WT mice infected with strains 25291 (blue) or 2447 (pink). (B and C) Bacterial load in the BM from WT, IFNγKO and iNOSKO mice infected with *M. avium* strain 25291 for 80 days. At each time-point of infection, groups were compared by two-tailed unpaired *t*-test, and were marked as: \*\*\**p* <0.001; **** *p* <0.0001. (D) Total number of BM cells per femur of non-infected WT mice (white), or after infection with *M. avium* strain 25291 (blue) or strain 2447 (pink) at 30, 60 and 80 dpi. The groups of infected mice, at the several time-points, were compared with the non-infected mice by ordinary one-way ANOVA followed by Dunnett’s multiple comparisons test, and were marked as: * *p* <0.05; ** *p* <0.01; comparisons between infected groups were performed by 2-way ANOVA followed by Tukey’s multiple comparisons test, and marked as: # *p* <0.05, ## *p* <0.01. Data represent the mean ± SD (A) or mean (B-D) from 5 to 8 mice per group, from one of two independent experiments. NI stands for non-infected.

To study the production of T cell precursor cells in the BM during infection by *M. avium* strain 25291, we analyzed the different BM cell populations by flow cytometry. Hematopoietic stem cells (HSC) possess the capacity of self-renewal, can give rise to all mature cell types found in the blood and are part of the LSK (Lineage^−^ Sca1^+^ cKit^+^) population ^44^ (see gating strategy, Fig. 2A). In comparison to non-infected mice, animals infected with *M. avium* strain 2447 had a transient increase in LSK cells percentage at 30 dpi, which returned to normal levels over time. In mice infected with highly virulent strain 25291, LSK cells percentage increase was sustained throughout the infection (Fig. 2B). IFNγ was shown to induce LSK cell expansion ^45^. Accordingly, upon IFNγKO mice infection with *M. avium* strain 25291, no alterations on LSK cells percentage at 80 dpi were observed (Fig. 2C).

**Figure 2.**
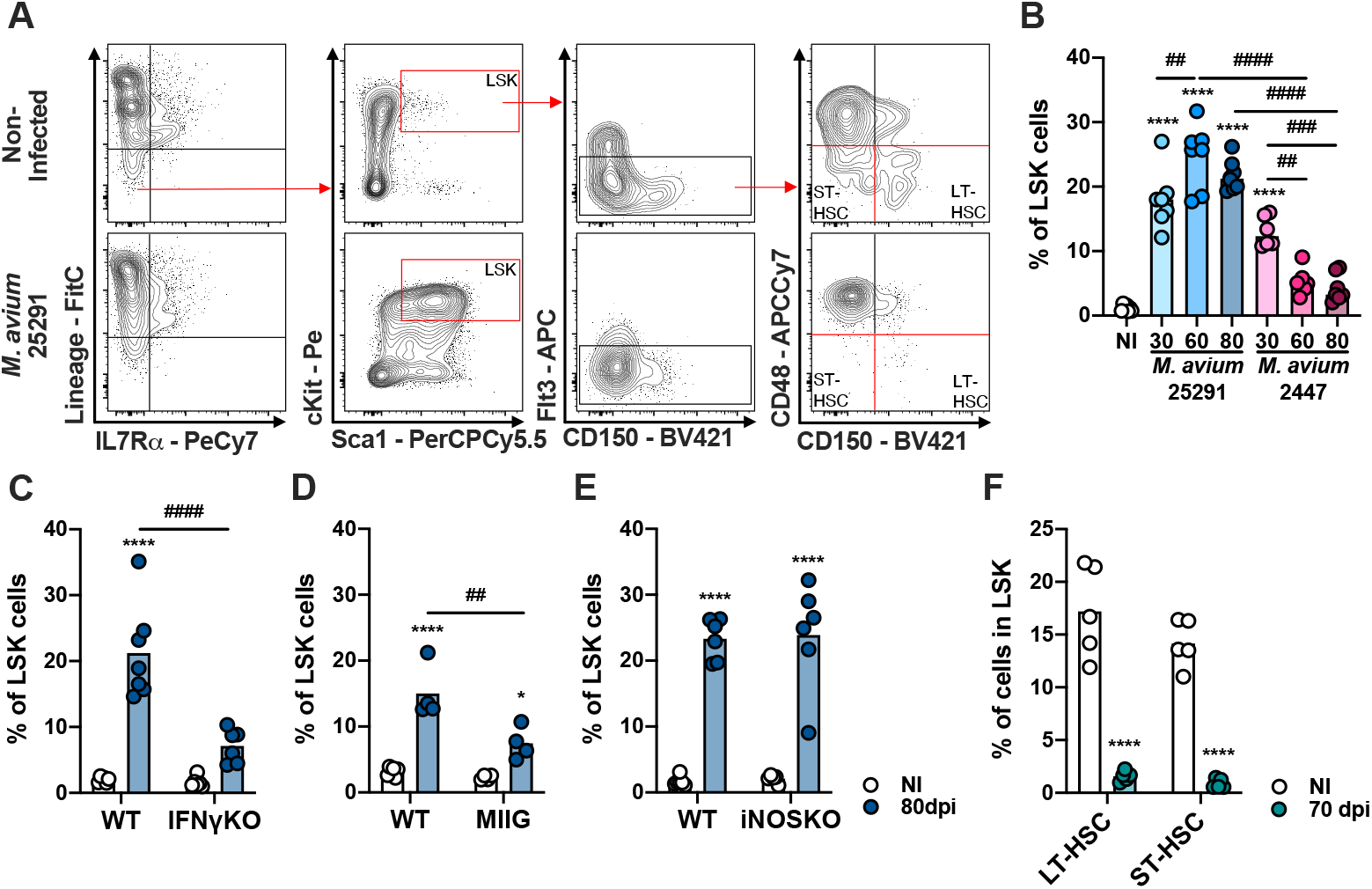
Mice infected with *M. avium* strain 25291 have higher percentage of LSK cells in an IFNγ-dependent and iNOS-independent manner. (A) Schematic representation of the gating used to identify LSK, LT-HSC and ST-HSC cell populations in the BM from non-infected WT mice (top), or infected for 70 days with *M. avium* 25291 (bottom). Total BM cells were previously selected eliminating doublets and debris. (B) Percentage of LSK cells from non-infected WT mice (white) or after infection with *M. avium* strain 25291 (blue) or strain 2447 (pink) at 30, 60 and 80 dpi. Comparisons between infected and non-infected mice were performed using ordinary one-way ANOVA followed by Dunnett’s multiple comparisons test, and marked as: **** *p* <0.0001; comparisons between all the infected groups were performed using a 2-way ANOVA followed by Tukey’s multiple comparisons test, and marked as: ## *p* <0.01, ### *p* <0.001, #### *p* <0.0001. (C, D and E) Percentage of LSK cells from non-infected (white) or infected for 80 days with *M. avium* 25291 (blue) of WT, IFNγKO, MIIG or iNOSKO mice. Groups were compared using 2-way ANOVA followed by Tukey’s multiple comparisons test; statistical differences between non-infected and infected mice were marked as * *p* <0.05 and **** *p* <0.0001; and between infected groups as ## *p* <0.01, #### *p* <0.0001. (F) Percentage of LT-HSC and ST-HSC cells in WT non-infected (white) or after 70 days of infection with *M. avium* 25291 (teal). Comparisons between infected and non-infected were performed by two-tailed unpaired *t*-test and marked as **** *p* <0.0001. Bars represent the mean from 4 to 8 mice per group from one of two independent experiments. NI stands for non-infected.

To understand if the increase in the percentage of LSK population was due to a direct effect of IFNγ, or through other mechanism dependent on Mϕ activation, we infected MIIG mice. At 80 dpi, MIIG mice showed increased LSK cells percentage when compared to non-infected mice. However, this increase was significantly smaller than the one observed on WT infected mice (Fig. 2D). As for iNOSKO mice, we show that the percentage of LSK cells increased upon infected with strain 25291 (80 dpi), as seen for WT mice (Fig. 2E). Together, these results show that the increase on LSK cells percentage during infection by *M. avium* strain 25291 is partially dependent on IFNγ-induced activation of Mϕ but not on iNOS production. Regarding HSC, we observed that infection by *M. avium* strain 25291 led to a dramatic decrease in the percentage of both long term (LT)-HSC and short term (ST)-HSC (Fig. 2F).

### Reduced proportion of LMPP and CLP cells in the BM is partially dependent on IFNγ but not iNOS

To our knowledge, there are two BM cell populations described to have the potential to differentiate into T cells *in vitro* and *in vivo*^46^: (1) the lymphoid-primed multipotent progenitors (LMPP), which is a subpopulation of LSK cells that express Flt3 (Lin^−^ IL-7Rα^−^ Sca1^+^ cKit^+^ Flt3^+^-gating strategy in Fig. 3A); and (2) the common lymphoid progenitors (CLP), which are Lin^−^ IL-7Rα^+^ Sca1^int^ cKit^int^ Flt3^+^ (gating strategy in Fig. 3F). In comparison to non-infected WT mice, the percentage of LMPP cells among LSK cells was decreased in mice infected with either *M. avium* strain 25291 or strain 2447 (Fig. 3B). The reduction in the percentage of LMPP suggests that the observed expansion of LSK cells cannot be attributed to an increase of LMPP cells. The reduction on the percentage of LMPP was not observed in infected IFNγKO nor in MIIG mice, but was preserved in iNOSKO mice (Fig. 3C, 3D, 3E). These results show that the decrease in the proportion of LMPP is mediated by the activation of Mϕ by IFNγ and independent of iNOS expression.

**Figure 3.**
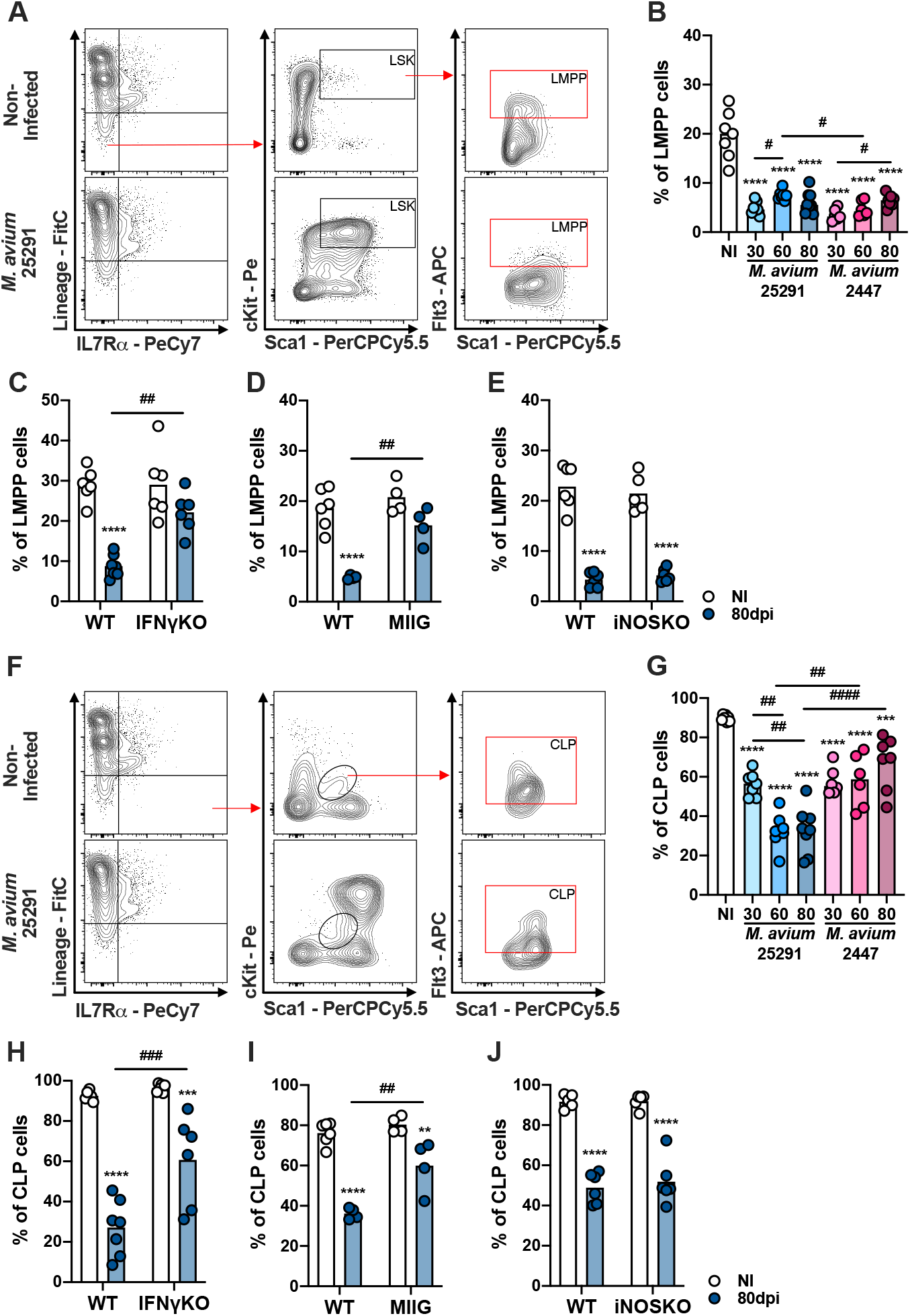
The reduction on the percentage of LMPP and CLP cells in mice infected with *M. avium* 25291 is dependent on IFNγ but not on iNOS. (A) Schematic representation of the gating used to select LMPP population in the BM from non-infected WT mice (top), or infected for 70 days with *M. avium* 25291 (bottom). Total BM cells were previously selected eliminating doublets and debris. (B) Percentage of LMPP cells from non-infected (white) WT mice or infected with *M. avium* strain 25291 (blue) or strain 2447 (pink) at 30, 60 and 80 dpi. (C, D and E) Percentage of LMPP cells from non-infected (white) or infected for 80 days with *M. avium* 25291 (blue), WT, IFNγKO, MIIG or iNOSKO mice. (F) Schematic representation of the gating used to select CLP population in the BM from non-infected WT mice (top), or infected for 70 days with *M. avium* 25291. Total BM cells were selected eliminating doublets and debris. (G) Percentage of CLP cells from non-infected WT mice (white), or infected with *M. avium* strain 25291 (blue) or strain 2447 (pink) at 30, 60 and 80 dpi. (H, I and J) Percentage of CLP cells from non-infected (white) or infected for 80 days with *M. avium* 25291 (blue), WT, IFNγKO, MIIG or iNOSKO mice. In all graphs, bars represent the mean from 4 to 8 mice per group, from one of two independent experiments. In B and G comparisons between infected and non-infected mice were evaluated by ordinary one-way ANOVA followed by Dunnett’s multiple comparisons test and marked as: *** *p* <0.001, **** *p* <0.0001; comparisons between infected groups were evaluated by 2-way ANOVA followed by Tukey’s multiple comparisons test and marked as: # *p* <0.05, ## *p* <0.01, #### *p* <0.0001. In C-E and H-J, comparisons were evaluated by 2-way ANOVA followed by Tukey’s multiple comparisons test and marked as ** *p* <0.01, *** *p* <0.001, **** *p* <0.0001 for comparisons between non-infected and infected, and as ## *p* < 0.01, ### *p* <0.001 for comparisons between infected groups. NI stands for non-infected.

We observed an obvious decrease in the percentage of CLP in WT mice infected with *M. avium* strain 25291, a change that is significantly smaller in mice infected with the low virulence strain 2447 (Fig. 3G). The decrease in the percentage of CLP upon infection with strain 25291 is partially dependent on IFNγ, since IFNγKO and MIIG mice also had lower percentages of CLP in comparison to non-infected mice, although significantly higher than observed in infected WT mice (Fig. 3H, 3I). iNOSKO mice infected with *M. avium* strain 25291 had a reduction in the percentage of CLP cells similar to WT infected mice (Fig. 3J). These results demonstrate that IFNγ and IFNγ signaling in Mϕ, but not the production of NO, are partially responsible for the alterations in CLP cells after *M. avium* strain 25291 infection.

Since the alterations on LMPP and CLP were not dependent on iNOS, this pathway was excluded as a potential mechanism responsible for *M. avium*-induced alterations on T cell BM precursors; still it might interact with other mechanisms yet to be identified. To assess which other molecules might be related to the alterations on BM cell populations, we analyzed the expression of previously described mediators associated with different models of BM alterations caused by infection/inflammation, specifically *Ifng,* glucocorticoid receptor (*Gr*), interleukine-6 (*Il6*) and tumor necrosis factor (*Tnf*). We observed that the BM of infected iNOSKO mice presents higher *Ifng* expression, and that the expression of *Il6* was increased in WT mice infected with strain 25291 when compared to non-infected mice, but not on infected iNOSKO mice (Sup. Fig. 1).

### Bone marrow cells from mice infected with *M. avium* 25291 poorly reconstitute thymi from RAGKO mice

Our results suggest that alterations in the BM precursors might be, at least in part, responsible for infection-induced thymic atrophy. To investigate this hypothesis, we transferred BM cells (lineage negative) from WT or IFNγKO mice, non-infected or infected for 70 days with *M. avium* strain 25291, into RAGKO recipients. Mice were sacrificed 4 weeks later. We observed that mice receiving BM cells from WT mice infected with *M. avium* 25291 had lower reconstitution of the recipient thymus than the ones with BM cells coming from non-infected mice or from IFNγKO infected mice (Fig. 4A left). When the four main thymocyte populations [double negative (DN; CD4^−^CD8^−^), double positive (DP; CD4^+^CD8^+^), CD4 single positive (CD4SP; CD4^+^CD8^−^) and CD8 single positive (CD8SP; CD4^−^CD8^+^)] were analyzed, we saw a significant decrease in the number of CD4SP and CD8SP thymocytes in mice that received BM cells from WT infected group. No differences were observed for mice that received BM cells from IFNγ infected group in comparison to non-infected (Fig. 4A right). In agreement, a lower number of total spleenocytes, and in particular of T cells (total and both CD4^+^ and CD8^+^), were recovered from mice receiving BM from infected WT mice, when compared to mice receiving BM cells from non-infected WT or IFNγ infected mice (Fig. 4B).

**Figure 4.**
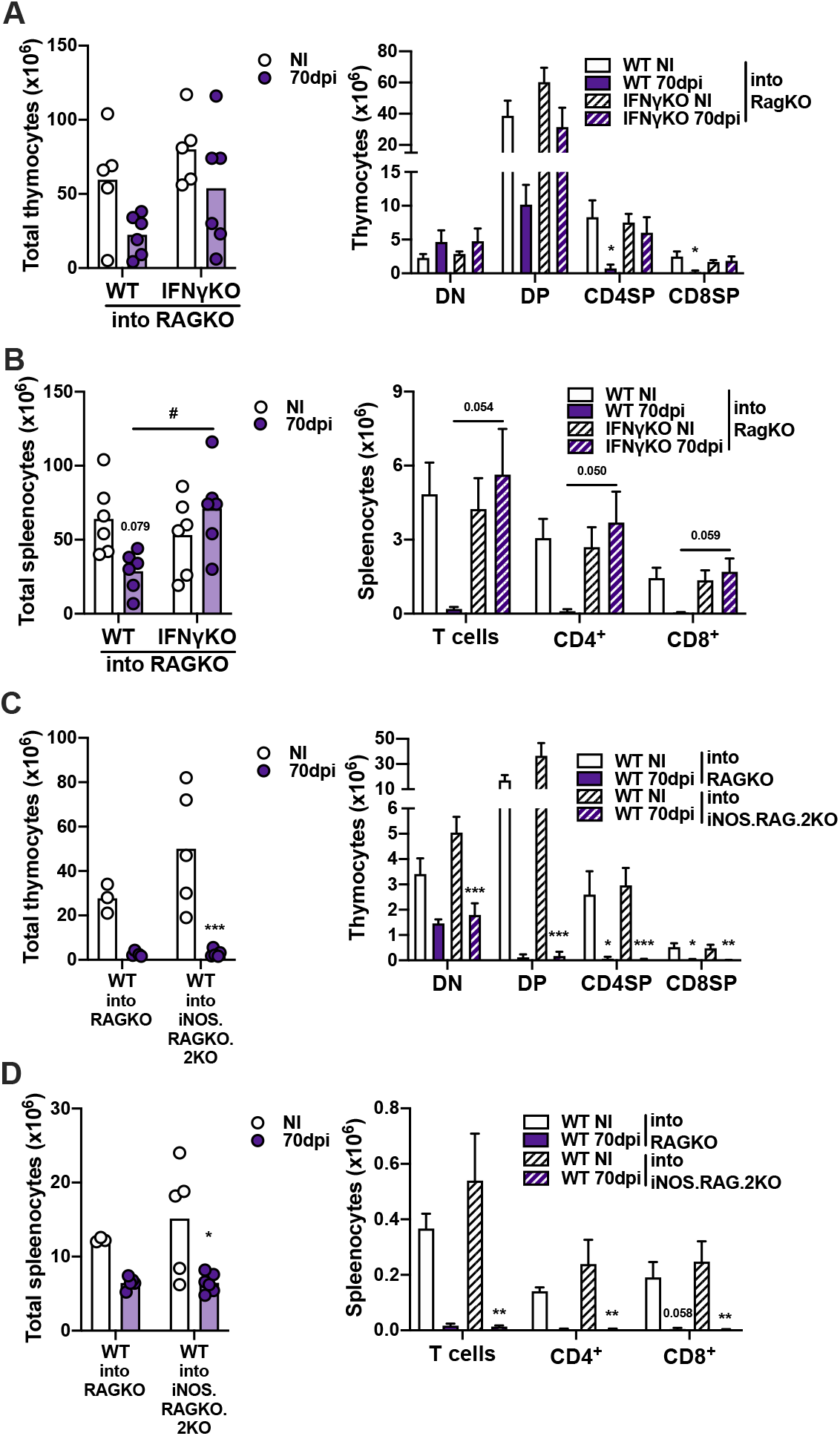
Inefficient reconstitution of thymi from RAGKO mice by BM cells from *M. avium* 25291 infected mice is IFNγ-dependent. (A-B) RAGKO mice were reconstituted with lineage negative BM cells from non-infected (white) or 70 days *M. avium* 25291 infected (purple) WT or IFNγKO mice. (A) Number of total thymocytes (left) and of the four main thymocyte populations (DN, DP, CD4SP and CD8SP). (B) Number of total spleenocytes (left) and of splenic total T cells and of CD4^+^ and CD8^+^ T cells (right). (C-D) RAGKO or iNOS.RAG.2KO mice were reconstituted with lineage negative BM cells from non-infected (white) or 70 days *M. avium* 25291 infected (purple) WT mice. (C) Number of total thymocytes (left) and of cells from the four main thymocyte populations (right). (D) Number of total spleenocytes (left) and of splenic T cells and its subpopulations (right). Bars represent the mean (left graphs) or the mean + SD (right graphs) from 3 to 6 mice per group from one of two independent experiments. Comparisons were performed using 2-way ANOVA followed by Tukey’s multiple comparisons test, and marked between infected and non-infected mice as * *p* <0.05, ** p <0.01, *** *p* <0.001; and between infected groups as # *p* <0.05. When *p*-values between [0.05; 0.10] (tendencies), their values are represented in the graphs. NI stands for non-infected.

To test if the reconstitution ability of BM precursor cells from WT infected mice was hampered by the NO production in the recipient thymus, we transferred BM cells (lineage negative) from non-infected or infected for 70 days with *M. avium* 25291 WT mice, into iNOS.RAG.2KO recipients. Irrespective of donor BM cells being from infected or non-infected mice, no differences between RAGKO and iNOS.RAG.2KO recipient mice were observed when comparing the reconstitution of the thymus (Fig. 4C), and the T cell pool in the spleen (Fig. 4D). These results indicate that, BM precursor cells from mice infected for 70 days present lower ability to reconstitute mice lacking T cells in a process dependent on the expression of IFNγ production but independent of NO production by the recipient thymic stromal cells. The data presented here support the hypothesis that infection-induced BM cells alterations play a relevant part on infection-induced premature thymic atrophy.

### iNOS expression is associated with a more inflammatory profile in the thymus after infection

The observation that iNOSKO mice present the same alterations on BM precursor cells after infection, but no premature thymic atrophy, suggests that IFNγ-induced BM alterations *per se* are not sufficient to cause premature thymic atrophy. So, we hypothesized that NO production is having an impact directly in the thymus. To investigate this, we evaluated the expression of inflammatory molecules in the thymus of WT and iNOSKO mice. We observed an increased expression of *Ifng* upon infection, both in the presence and absence of iNOS expression. (Fig. 5A). *Gr* gene expression was not altered in the thymus after infection with *M. avium* strain 25291 (Fig. 5B). Additionally, WT, but not iNOSKO infected mice, presented increased corticosterone levels in the serum (Fig. 5C). The absence of NO production, together with low production of corticosterone may contribute to the nonexistence of *M. avium*-induced premature thymic atrophy in iNOSKO mice, even presenting alterations in BM T cell precursor cells.

**Figure 5.**
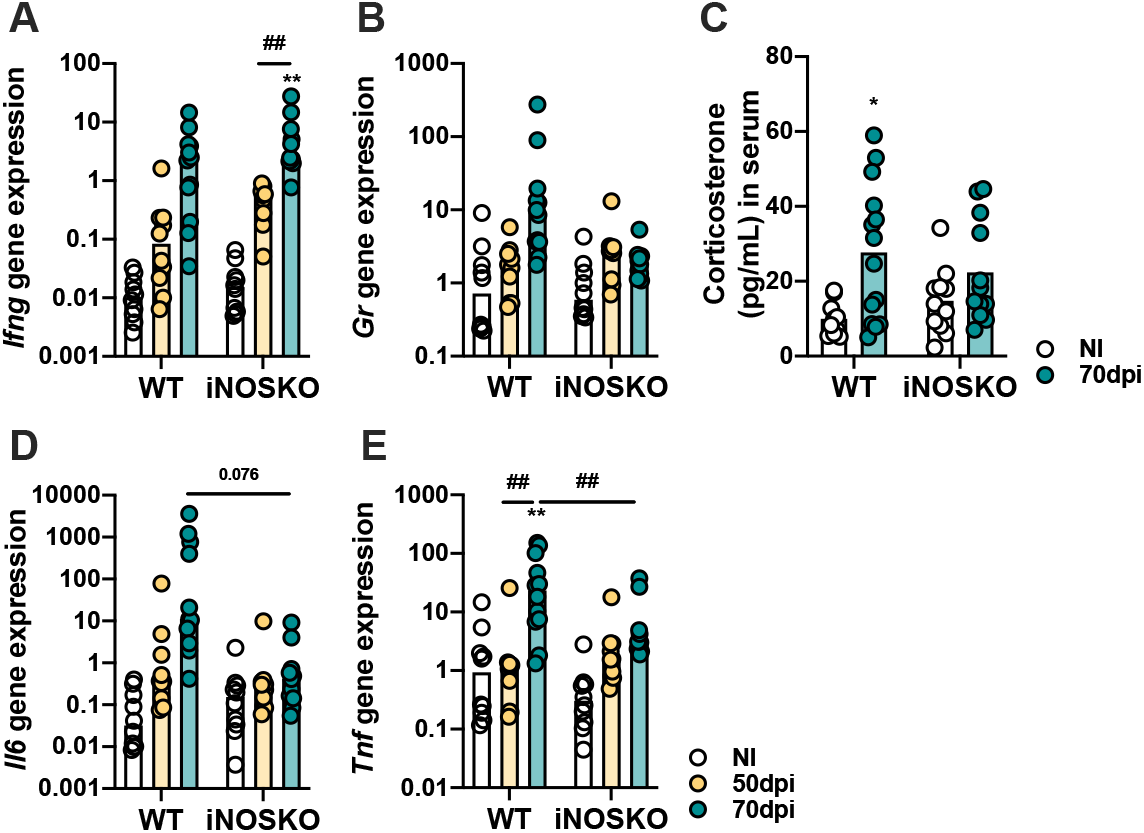
*M. avium* strain 25291 infected iNOSKO mice present a milder inflammatory profile in the thymus than WT infected mice. RNA expression levels of *Ifng* (A), *Gr* (B), *Il6* (D) and *Tnf* (E) in the thymus of WT and iNOSKO mice non-infected (white) or infected with *M. avium* strain 25291 for 50 (yellow) or 70 (teal) days. (C) Basal corticosterone levels in the serum of WT and iNOSKO mice non-infected (white) or infected with *M. avium* strain 25291 for 70 days (teal). Bars represent the median for all graphs except for C where mean is represented, from 10 to 12 mice per group from two pooled independent experiments. Statistically significant differences were accessed by 2-way ANOVA followed by Tukey’s multiple comparisons test and marked between non-infected and infected groups as: * *p* <0.05, ** *p* <0.01; and between infected groups as: ## *p* <0.01. NI stands for non-infected.

Both IL-6 and TNF have been associated with premature thymic atrophy. Here we observed that, in comparison to infected iNOSKO mice, WT infected mice present a tendency for increased *Il6* expression, and a significant increase on *Tnf* expression in the thymus (Fig. 5D, 5E). These data indicate that NO production is associated with increased inflammation in the thymus after infection, and that this might negatively impact on thymocyte differentiation and/or potentially be a mechanism of thymocyte death.

### Thymic stroma from *M. avium* 25291 infected mice is unable to support optimal T cell differentiation

To determine if alterations in the thymic stroma could contribute to infection-induced thymic atrophy, we investigated the ability of the thymus from infected animals to support thymocyte differentiation when BM precursor cells from non-infected mice are provided. For this purpose, atrophied thymi from infected WT mice (CD45.2) and from uninfected RAGKO mice (CD45.2), were transplanted under the kidney capsule of uninfected WT CD45.1 mice (one thymus on each kidney of the same mouse; Fig. 6A). Four weeks after transplant mice were euthanized and the four main thymocyte populations within CD45.1^+^ cells were analyzed in the transplanted thymi and in the endogenous thymi of recipient mice (Fig. 6B). We observed that thymocyte populations in the WT infected thymi were completely altered in comparison to RAGKO thymi, which were comparable to the ones observed in the endogenous thymi (Fig. 6C). Furthermore, the number of cells from each thymocyte population was higher in non-infected RAGKO thymi than in thymi from infected WT mice (Fig. 6D). These results show that thymic stroma from WT mice infected with *M. avium* 25291 have impaired ability to support thymocyte differentiation.

**Figure 6.**
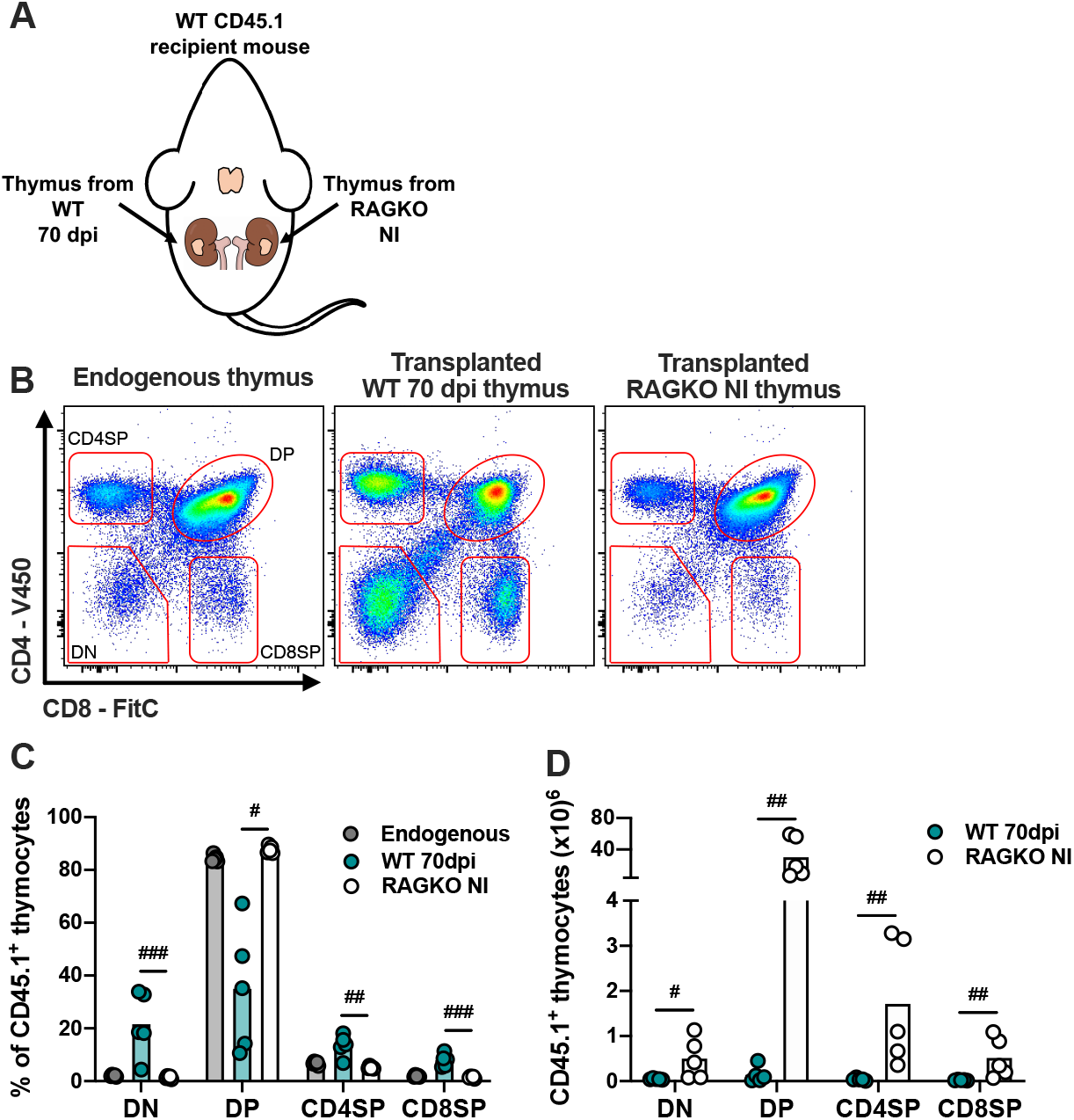
Thymic stroma from WT mice infected with *M. avium* 25291 is unable to properly support T cell differentiation. (A) Schematic representation of thymi transplant from *M. avium* infected WT mice (70 dpi) and non-infected RAGKO mice (both CD45.2) under the kidney capsule of WT CD45.1 recipient mice. (B) Representative plots of the CD45.1^+^ four main thymocyte populations (DN, DP, CD4SP and CD8SP) from WT CD45.1 recipient mice endogenous thymi, and from transplanted WT infected and RAGKO non-infected thymi. (C) Percentage and (D) number of cells from the four main thymocyte populations from endogenous thymi of WT CD45.1 recipient mice (grey), or from transplanted thymi from 70 days *M. avium* 25291 infected WT mice (teal) or non-infected RAGKO mice (white). Data represent the mean from five mice per group from one of two independent experiments. Comparisons between WT 70dpi and non-infected RAGKO were performed by 2-tailed ratio paired t test and marked as: # *p* <0.05, ## *p* <0.01, ### *p* <0.001. NI stands for non-infected.

### Increased thymocyte death in infected mice is independent of caspase-3 activation

To further dissect the mechanisms leading to infection-induced thymic atrophy we investigated thymocytes’ death by analyzing the incorporation of Propidium iodide (Pi) and Annexin V stain in WT mice infected with *M. avium* strain 25291 or strain 2447 at 30, 60 and 70 dpi (Fig. 7A). Infection by both *M. avium* strains led to a decrease on the percentage of viable cells (Annexin V^−^ Pi^−^) and an increase on the percentage of apoptotic cells (Annexin V^+^ Pi^−/low^; Fig. 7A). However, in mice infected with the high virulence strain 25291, these alterations occurred earlier and were greater than the ones in mice infected with the low virulence strain 2447. Increased percentage of thymocytes undergoing necrosis/late apoptosis (Annexin V^+^ Pi^high^) was observed at 70 dpi with *M. avium* strain 25291, but not with strain 2447 (Fig. 7A).

**Figure 7.**
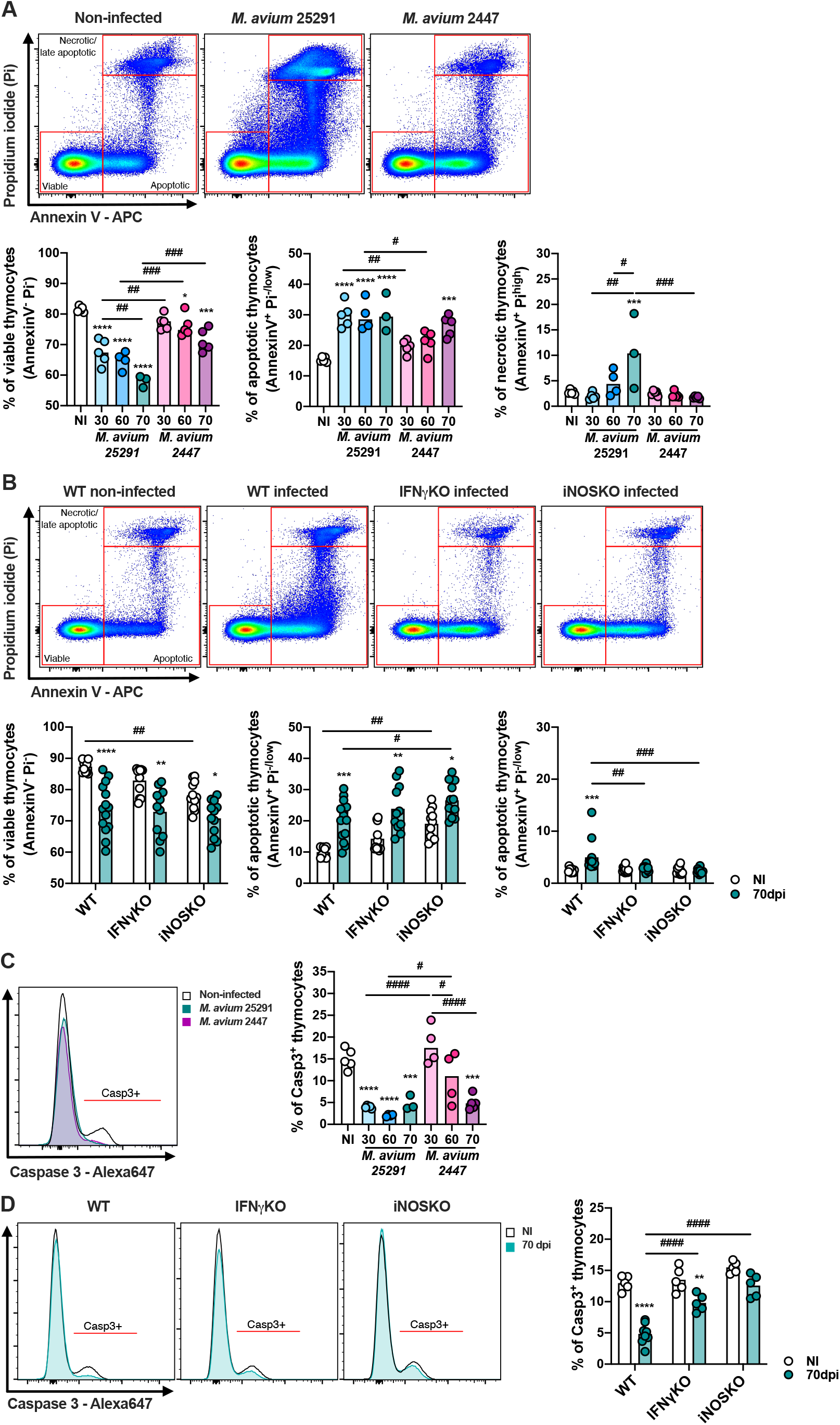
The percentage of viable cells and of cells positive for active caspase-3 decrease after infection. (A) Representation of the gating and plotting of viable (left – AnnexinV^−^ Pi^−^), apoptotic (center – AnnexinV^+^ Pi^−/low^) and necrotic/late apoptotic (right – AnnexinV^+^ Pi^high^) thymocytes from non-infected WT mice (white) or infected for 30, 60 or 70 days with *M. avium* 25291 (blue) or *M. avium* 2447 (pink). (B) Representation of the gating and plotting of select viable (left – AnnexinV^−^ Pi^−^), apoptotic (center –AnnexinV^+^ Pi^−/low^) and necrotic/late apoptotic (right –AnnexinV^+^ Pi^high^) thymocytes from WT, IFNγKO or iNOSKO mice non-infected (white) or infected for 70 days with *M. avium* 25291 (teal). (C) Representative histogram and plotting of activated caspase 3 positive thymocytes from non-infected WT mice (white) or infected for 30, 60 or 70 days with *M. avium* 25291 (blue) or *M. avium* 2447 (pink). (D) Representative histogram and plotting of caspase 3 positive thymocytes from WT, IFNγKO or iNOSKO mice non-infected (white) or infected for 70 days with *M. avium* 25291 (teal). Total thymocytes were previously selected eliminating doublets and debris. Bars represent the mean from: 3 to 5 mice per group from one experiment for A and C; 9 to 14 mice per group from two pooled independent experiments for B and D. In A and C comparisons between infected and non-infected mice were evaluated by ordinary one-way ANOVA followed by Dunnett’s multiple comparisons test and marked as: * *p* <0.05, *** *p* <0.001, **** *p* <0.0001; and comparisons between infected groups were evaluated by 2-way ANOVA followed by Tukey’s multiple comparisons test and marked as: # *p* <0.05, ## *p* <0.01, ### *p* <0.001, #### *p* <0.0001. In B and D, comparisons were performed by 2-way ANOVA followed by Tukey’s multiple comparisons test, and marked as * *p* <0.05, ** *p* <0.01, *** *p* <0.001, **** *p* <0.0001 for differences between non-infected and infected, and as # *p* <0.05, ## *p* <0.01, ### *p* <0.001, #### *p* <0.0001 for differences between infected groups. NI stands for non-infected. Casp3 stands for active caspase 3.

To understand if these alterations depend on IFNγ and/or NO production, we analyzed thymocyte cell death on thymi from IFNγKO and iNOSKO mice at 70dpi. The decrease on the percentage of viable thymocytes, and the increase on the percentage of apoptotic thymocytes was also observed in IFNγKO and iNOSKO mice (Fig. 7B), indicating that these molecules do not play an essential role in this process. However, both IFNγKO and iNOSKO infected mice showed no alterations on the percentage of necrotic/late apoptotic thymocytes in comparison to the non-infected peers (Fig. 7B). In respect to cell death in the four main thymocyte populations, DP, CD4SP and CD8SP were the ones most affected after infection, in an IFNγ and NO independent manner (Sup. Fig. 2). In infected WT mice, DP thymocytes were the only population presenting higher percentage of cells in necrosis/late apoptosis, a difference not present in IFNγKO and iNOSKO infected mice, as observed in total thymocytes (Sup. Fig. 2B). This indicates that DP thymocytes’ necrosis/late apoptosis induced by *M. avium* strain 25291 infection is dependent on IFNγ and NO.

We observed that infection with both *M. avium* strains lead to decreased percentages of activated caspase 3 positive cells. On mice infected with strain 25291, this decrease is evident as soon as 30 dpi, and sustained, while this was only observed in mice infected with strain 2447 at 70 dpi (Fig. 7C). Upon infection with strain 25291, IFNγKO mice showed decreased percentages of activated caspase 3 positive cells, though it did not reach the values observed for WT infected mice, and iNOSKO mice presented no alterations (Fig. 7D). These results show that the increased apoptosis on thymocytes after infection is independent of caspase 3 activation. And that the decrease in active caspase 3 positive cells is partially dependent of IFNγ and dependent of iNOS.

## DISCUSSION

Infection by *M. avium* strain 25291 results in severe thymic atrophy, a process dependent on the synergy between GC and NO produced by IFNγ activated Mϕ ^10^. Here we explored the mechanisms affecting the thymus and the BM T cell precursors, that could be responsible for this infection-induced thymic atrophy. This study shows that the process leading to *M. avium*-induced thymic atrophy results from a combination of the effect of IFNγ and NO at distinct steps of T cell differentiation, affecting T cell precursors in the BM, thymocytes and thymic stroma.

Here we show an increase in the percentage of LSK cells in the BM dependent on IFNγ, after infection with *M. avium* strain 25291. Our results are consistent with previous reports describing LSK expansion during infection by *M. tuberculosis*, *M. avium* and by Vaccinia virus, ^29,30,35,45^. To our knowledge, we show for the first time, that the expansion of LSK cells during *M. avium* infection is at least partially dependent on the signaling of IFNγ on the Mϕ, as MIIG mice (that have IFNγ production but no signaling of this cytokine on the CD68^+^ cells) present an increase on the percentage of LSK cells at 80 dpi, that is significantly lower than the one observed in infected WT mice. The expansion of LSK cells during *M. tuberculosis* infection was previously associated with two cytokines produced by Mϕ, TNF and IL-6 ^29^. In agreement we also observed an increase on the RNA expression levels of these two cytokines in the BM after *M. avium* strain 25291 infection, strengthening the role that the activation of Mϕ is playing. Within the LSK population, we observed a reduction on the percentage of the LT-HSC and of ST-HSC. HSC are very susceptible to the surrounding microenvironment and HSC alterations associated with infection and/or inflammation were already described. These alterations include reduction of the HSC pool and increased or decreased proliferation of this BM population ^30,47–52^. During chronic inflammation (or chronic IFNγ signaling) - which is the case after *M. avium* strain 25291 infection - there is impairment on the self-renewal capacity of HSC what can culminate in the reduction of this population ^47^; this might explain the reduction on the percentage of HSC that we observed in our model.

Regarding T cell precursor cells in the BM, CLP and LMPP, here we show that infection by *M. avium* induces a decrease in the percentage of LMPP cells that is dependent on the activation of Mϕ by IFNγ, and independent of iNOS. Kong *et al*., using a model of sepsis, associated a decrease in the percentage of LMPP with premature thymic atrophy ^39^, which is in accordance with our results. We also observed that, CLP cells were also reduced in percentage after *M. avium* infection, in a process partially dependent on IFNγ and independent on iNOS. These observations support the reduction of CLP cells observed by others during a thymic atrophy model of sepsis, and after *P. chabaudi* infection, though no association with thymic atrophy was mentioned in those studies ^36,39^. On the other hand, other infections, such as by Vaccinia virus, were shown to induce an increase of CLP cells in the BM ^35^, indicating that alterations in this population are infection specific and have not always been associated with premature thymic atrophy.

Unlike BM cells from non-infected WT mice or from IFNγKO infected mice, BM cells from *M. avium* infected WT mice presented impaired ability to reconstitute the thymus and the periphery of RAGKO mice. This result confirms that during infection by *M. avium* 25291, changes in the BM play a role in premature thymic atrophy. To our knowledge, there is only one report associating defects in the BM with premature thymic atrophy. This study showed that sepsis-induced thymic atrophy is associated with a dramatic decrease on the early thymic precursors (ETP) as a consequence of impaired migration of progenitors from the BM to the thymus, and of inability of lymphoid lineage commitment ^39^. Here we show a defect on the differentiation of precursor cells from the lymphoid lineage (LMPP and CLP); one other possibility is that IFNγ and/or NO might be down-regulating the expression of chemokine receptors responsible for the migration of T cell precursors from the BM to the thymus (such as CCR7, CCR9 and PSGL1), as it was shown in sepsis-induced thymic atrophy by others ^39^. Even though the BM presents severe alterations during *M. avium* strain 25291 infection, this does not seem to be the only causative mechanism of thymic atrophy. T cell differentiation only occurs in the thymic microenvironment, which is provided by thymic stromal cells. By performing thymic transplant under the kidney capsule, here we show that the stroma from *M. avium*-atrophied thymus presents an impairment in supporting the differentiation of new T cells when provided with BM precursors from non-infected WT mice, which are able to give rise to T cells in thymic stroma of RAGKO mice. Whether this impairment is related with diminished capacity to attract BM precursors, or due to limitations on supporting differentiation itself is unknown. However, on a model of thymic atrophy during pregnancy, thymic stromal cells had limited capacity to produce chemokines essential to the homing of T cell precursors to the thymus, such as CCL25, CXCL12, CCL21 and CCL19 ^53^. This hypothesis is consistent with our previous observation of a reduction on the most immature thymocytes (ETPs; T cell precursor cells that just entered the thymus) after infection with *M. avium* strain 25291 ^10^. It is also possible that infection of stromal cells leads to their degeneration and consequent thymocyte decline, as described during HIV infection ^54–56^.

Despite changes on thymic microenvironment during infection could induce thymocyte death, this process can also be caused by other mediators/mechanisms. Increased levels of GC, IFNγ, TNF and NO production were associated with thymocyte apoptosis during infection, or in other conditions ^25–28,57–59^. Here we show a decrease on the percentage of viable (Annexin V^−^ Pi^−^) thymocytes after infection, that is independent of both IFNγ and iNOS. During T cell differentiation, all the non-positively selected thymocytes die by programmed cell death. Since around 90% of the thymocytes die, the thymus became a specialized organ for dead cell clearance ^60^. This makes it challenging to study cell death in the thymus since dead cells are cleared very quickly and cannot be tracked, and may explain why only a slight decrease on the percentage of viable thymocytes after infection is observed. Still, the increased thymocyte death is consistent with most infection-induced thymic atrophy models, mostly *via* thymocyte apoptosis ^19,61–68^. We show increased *Tnf* RNA expression in the thymus after infection with *M. avium* strain 25291, which is a cytokine that may mediate thymocyte apoptosis, as reported in other models ^26,57,58^. However, the observed increase in thymocyte apoptosis was also observed in WT mice infected with *M. avium* strain 2447, and in IFNγKO and iNOSKO mice infected with strain 25291, and none of these models showed infection-induced thymic atrophy^10^. These observations imply that apoptosis, though it might be participating, should not be the sole mechanism responsible for *M. avium*-induced thymic atrophy. Additionally, we observed a decrease on the percentage of thymocytes positive for activated caspase 3. This means that though thymocytes are dying by apoptosis, this is not mediated by the activation of caspase 3, but by some other mechanism(s). However, as caspase 3 is fundamental for T cell differentiation in the thymus ^69^, the reduction on the activation of this enzyme suggests that other unexplored parameters of thymocytes’ differentiation are affected during infection, in an iNOS dependent way.

WT infected mice also show an increase on the percentage of necrotic/late apoptotic thymocytes, which was not shown for IFNγKO and iNOSKO infected mice. This increase was only observed at very late time points (70 dpi), and thymic atrophy is already evident earlier after infection ^10^, indicating that this mechanism is not the driver of *M. avium*-induced thymic atrophy, but perhaps a consequence.

As conclusion, here we show that *M. avium*-induced premature thymic atrophy is not the result of a single mechanism but of the association of several events: IFNγ-associated alterations on the T cell precursor cells in the BM, and on its ability to migrate to the thymus and/or differentiate into new T cells; reduced ability of thymic stromal cells to recruit T cell precursors and/or sustain T cell differentiation; and alterations on thymocyte apoptosis independent of both IFNγ and iNOS. The role of NO in the thymus during *M. avium*-induced thymic atrophy seems to be restricted to the alterations on caspase 3 activation in thymocytes, and on the inflammatory profile observed in this organ. Altogether our data suggest that, after infection with *M. avium* strain 25291, IFNγ is detrimental to BM T cell precursors, the ones that manage to reach the thymus encounter a harsh environment, resulting from NO-dependent inflammation, which renders most of the precursors unable to differentiate, and/or die, culminating in premature thymic atrophy.

## AUTHORS CONTRIBUTION

PB-S, MC-N and RA conceptualized, designed and supervised the study; PB-S performed the experiments and processed the samples, analyzed the data, prepared the figures, performed the statistical analysis and drafted the first version of the manuscript; RM-M, CN, SR, CS-M, MB and DSC collected and processed samples; MC-N, RA and SMB assembled the funding for experiments; and all authors discussed the results and contributed to the final version of the manuscript.

## DATA AVAILABILITY

The data that support the findings of this study are available from the corresponding author upon request.

