## Supplemental Figures for "IFNγ and iNOS-mediated alterations in the bone marrow and thymus and its impact on *Mycobacterium avium*-induced thymic atrophy"

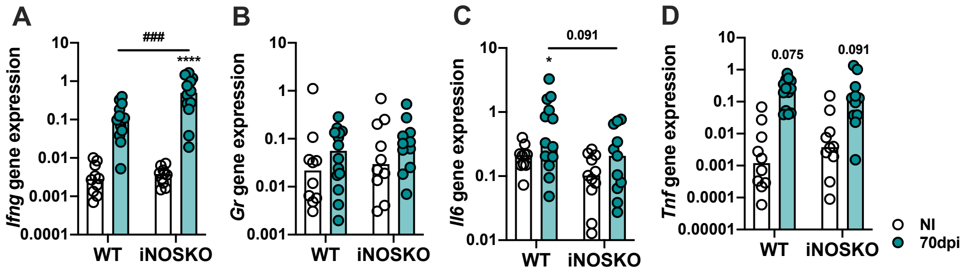


**Supplementary Figure 1. *M. avium* strain 25291 infected iNOSKO mice present higher *Ifng* and lower *Il6* expression in the BM.** RNA expression levels of *Ifng* (A), *Gr* (B), *Il6* (C) and *Tnf* (D) in the BM of WT and iNOSKO mice non-infected (white) or infected with *M. avium* strain 25291 for 70 days (teal). Bars represent the median from 10 to 14 mice per group from two pooled independent experiments. Comparisons were performed by 2-way ANOVA followed by Tukey's multiple comparisons test, and marked between non-infected and infected groups as: * $p$ <0.05, **** $p$ <0.0001; and between infected groups as ### $p$ <0.001. NI stands for non-infected.

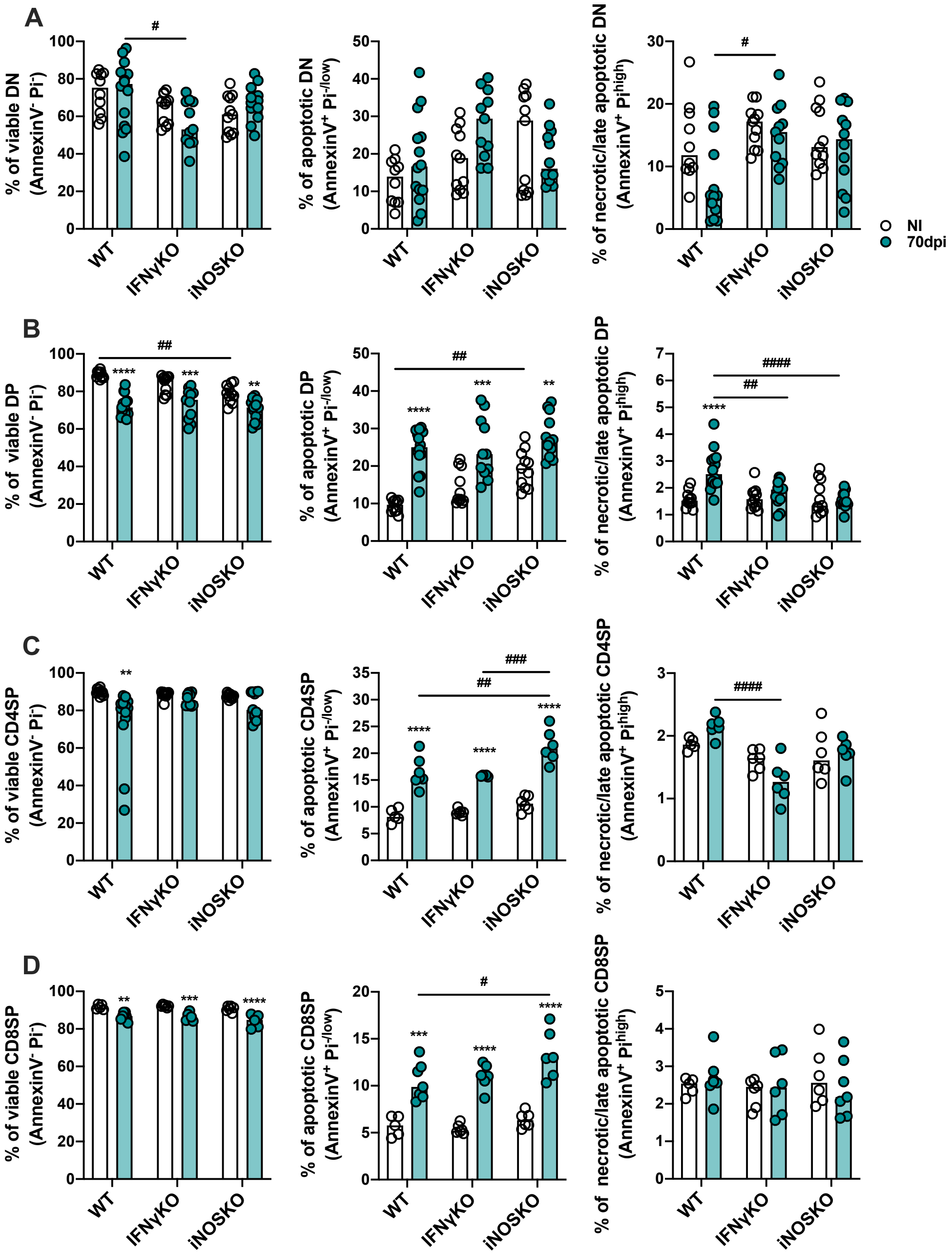


**Supplementary Figure 2. Decreased thymocyte viability is more evident in DP and SP thymocyte populations.** Viable (left – AnnexinV$^-$ Pi$^-$), apoptotic (center – AnnexinV$^+$ Pi$^{-/low}$) and necrotic/late apoptotic (right – AnnexinV$^+$ Pi$^{high}$), DN (A), DP (B), CD4SP (C) and CD8SP (D) thymocytes from WT, IFNγKO or iNOSKO mice non-infected (white) or infected for 70 days with *M. avium* 25291 (teal). Bars represent the mean from 9 to 14 mice per group from two independent experiments plotted together. Statistically significant differences were accessed by 2-way ANOVA followed by Tukey's multiple comparisons test and marked as ** $p$ <0.01, *** $p$ <0.001, **** $p$ <0.0001 between non-infected and infected, and as # $p$ <0.05, ## $p$ <0.01, ### $p$ <0.001, #### $p$ <0.0001 between infected groups. NI stands for non-infected.
